## Supplemental Material for "4-aminoquinolines block heme iron reactivity and interfere with artemisinin action"

### Figure 1- figure supplement 1

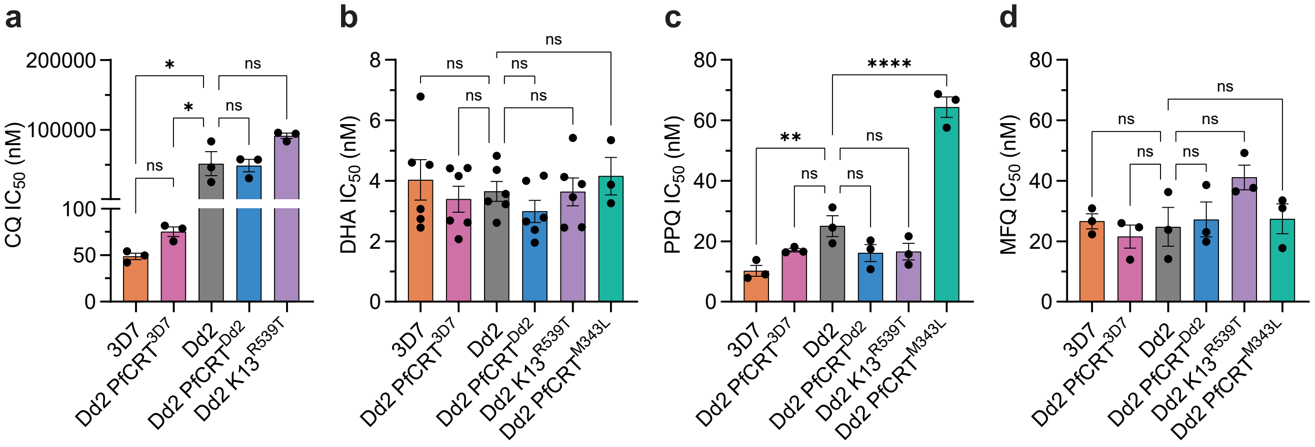


**Figure 1- figure supplement 1. CQ, DHA, PPQ, and MFQ sensitivity of different parasites.** Shown are mean (**a**) CQ, (**b**) DHA, (**c**) PPQ, and (**d**) MFQ trophozoite stage IC50 values ± SEM obtained as part of independent isobologram replicates (see Figure 1). For (**b**) DHA, IC50 values from CQ-DHA and PPQ-DHA isobolograms are shown. Statistical significance between Dd2 vs all other parasites was determined using a one-way ANOVA with a Dunnett's test for multiple comparisons. ****$p < 0.0001$; **$p < 0.01$; *$p < 0.05$; ns = not significant.

### Figure 2- figure supplement 1

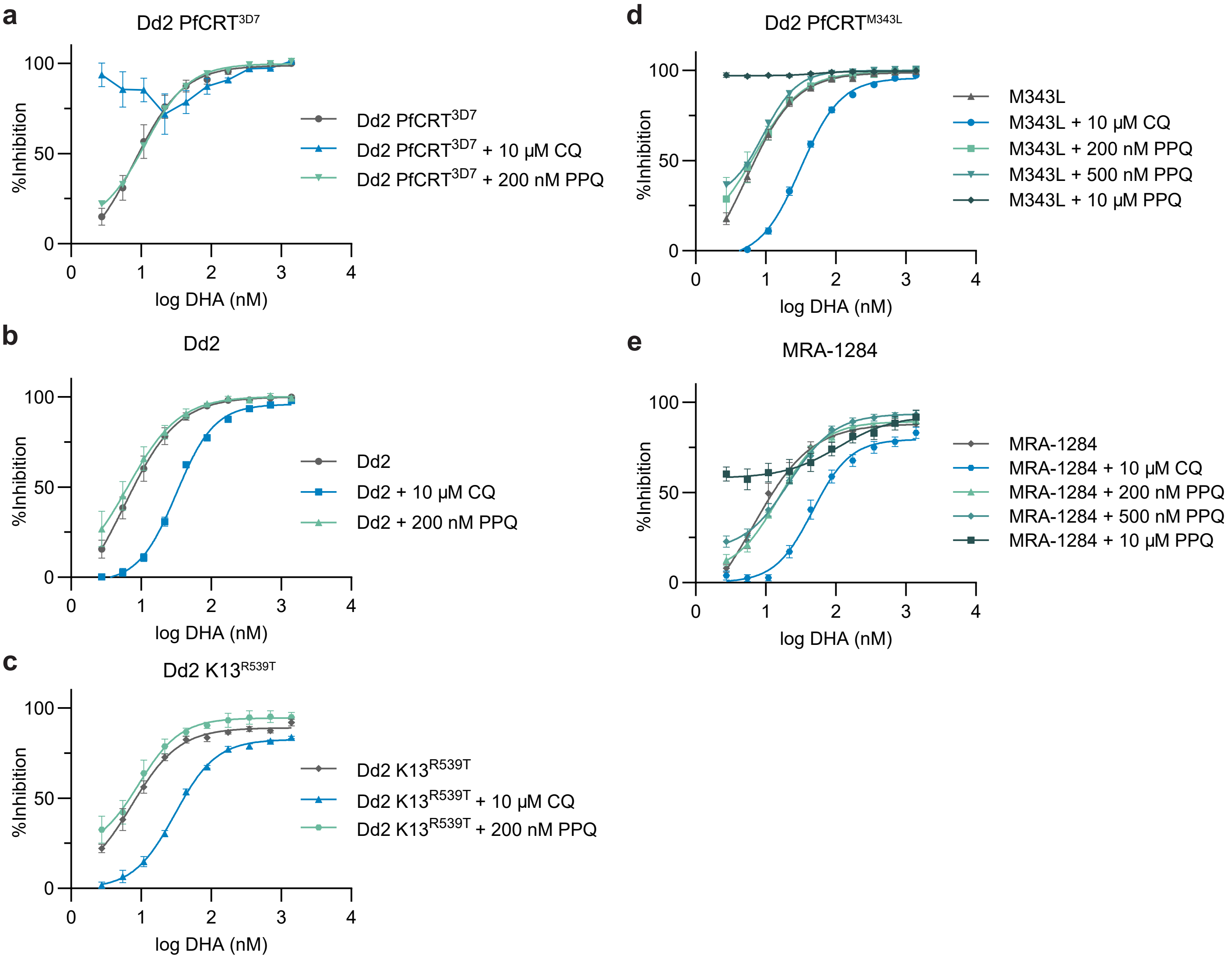


**Figure 2- figure supplement 1. Early ring stage dose-response curves.** DHA dose response assays on 0-3 hpi ring stage (**a**) Dd2 PfCRT$^{3D7}$, (**b**) Dd2, and (**c**) Dd2 K13$^{R539T}$ parasites were performed with DHA alone, or DHA with the addition of 10 µM CQ, or 200 nM PPQ. For the PPQ-resistant parasites (**d**) Dd2 PfCRT$^{M343L}$ and (**e**) MRA-1284, additional PPQ concentrations of 500 nM and 10 µM were also tested. Shown are dose response curves with mean % inhibition ± SEM from at least three independent replicates.

**Figure 3- figure supplement 1**

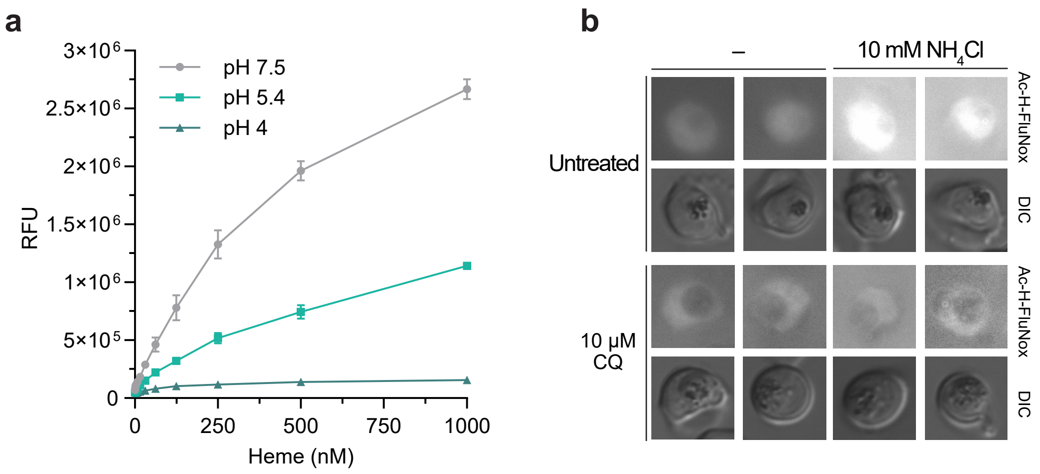


**Figure 3- figure supplement 1. Effect of pH on H-FluNox and Ac-H-FluNox activity.**

(**a**) H-FluNox was incubated with increasing concentrations of heme at pH 7.5, 5.4, and 4. Shown are mean relative fluorescence units (RFU) ± SD from two independent replicates performed in technical duplicate. (**b**) Dd2 trophozoites were treated with 10 µM CQ or mock treated for 5.5 h. Parasites were then co-incubated with Ac-H-FluNox and 10 mM ammonium chloride to visualize free heme and alkalinize the digestive vacuole, respectively. Shown are two representative images per treatment condition. Note that exposure times were optimized for different treatment conditions to better visualize the digestive vacuole.

### Figure 3- figure supplement 2

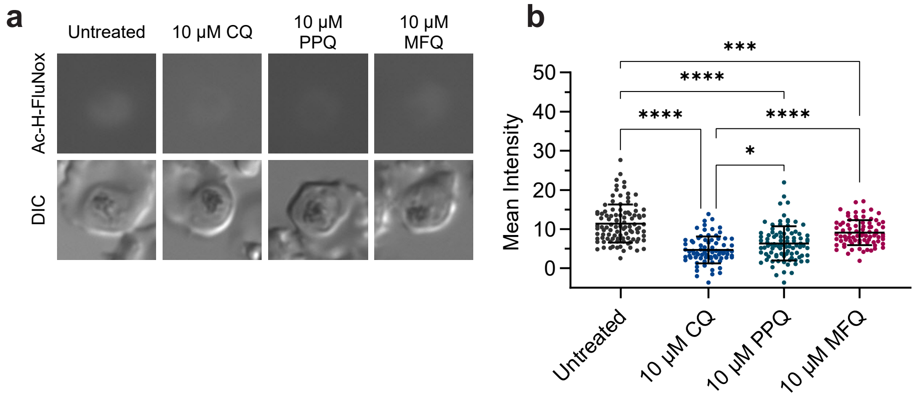


**Figure 3- figure supplement 2. Equimolar comparison of CQ, PPQ, and MFQ on Ac-H-FluNox fluorescence.** Dd2 trophozoites were treated with 10 μM CQ, 10 uM PPQ, 10 uM MFQ, or mock treated for 1 h. Parasites were then incubated with Ac-H-FluNox to visualize and quantify "active" heme. Shown are (**a**) representative images and (**b**) mean fluorescence intensity ± SD of at least 75 parasites from three independent drug treatments. Statistical significance was assessed using a one-way ANOVA with a Turkey's test for multiple comparisons.

**Figure 3- figure supplement 3**

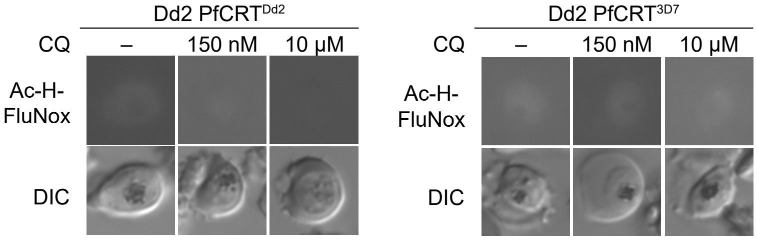


**Figure 3- figure supplement 3. Representative images of Dd2 PfCRT$^{Dd2}$ and Dd2 PfCRT$^{3D7}$ Ac-H-FluNox fluorescence.** Dd2 PfCRT$^{Dd2}$ and Dd2 PfCRT$^{3D7}$ trophozoites were mock treated or treated with 150 nM CQ or 10 µM CQ for 5.5 h. Parasites were then incubated with 10 µM Ac-H-FluNox (the cell permeable analog of H-FluNox) to visualize and quantify "active" heme. Shown are representative images relating to Figure 3f.

**Figure 3- figure supplement 4**

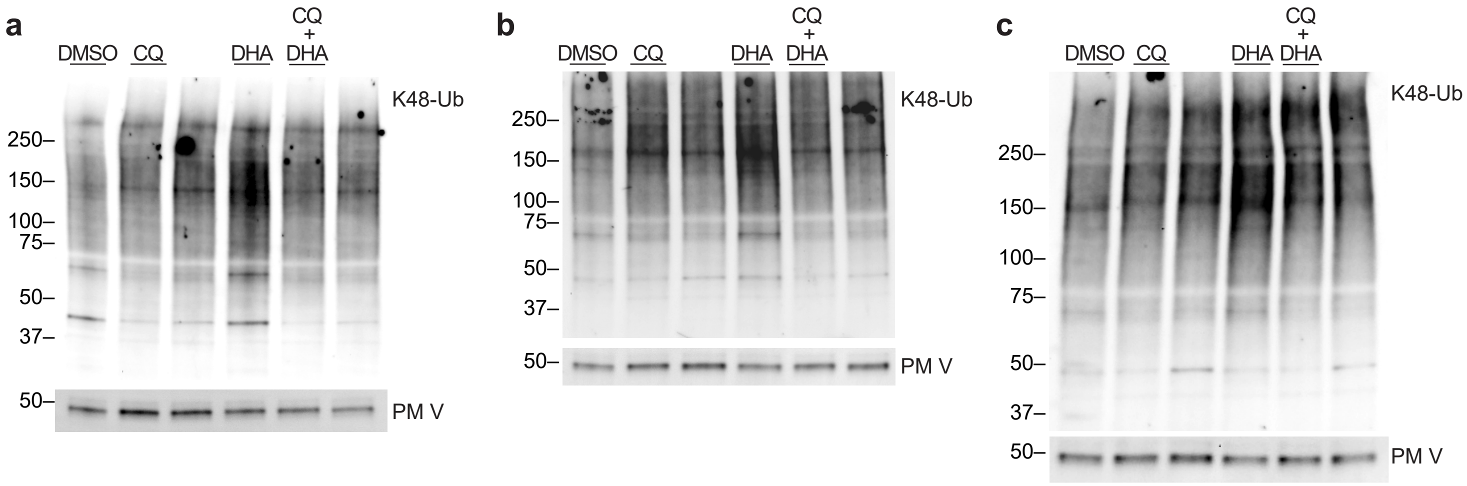


**Figure 3- figure supplement 4. Additional K48-ubiquitin western blots.**

Dd2 trophozoites were treated with 10 μM CQ, 25 nM DHA, 10 μM CQ + 25 nM DHA for 6 h. Parasite lysates were subjected to western blot and probed with anti-K48-linked ubiquitin (K48-Ub) antibodies or anti-plasmepsin V (PMV) antibodies. Blots from three independent replicates are shown. Note that (**a**) is an uncropped version of the blot in Figure 3g**.**

**Figure 3- source data 1**

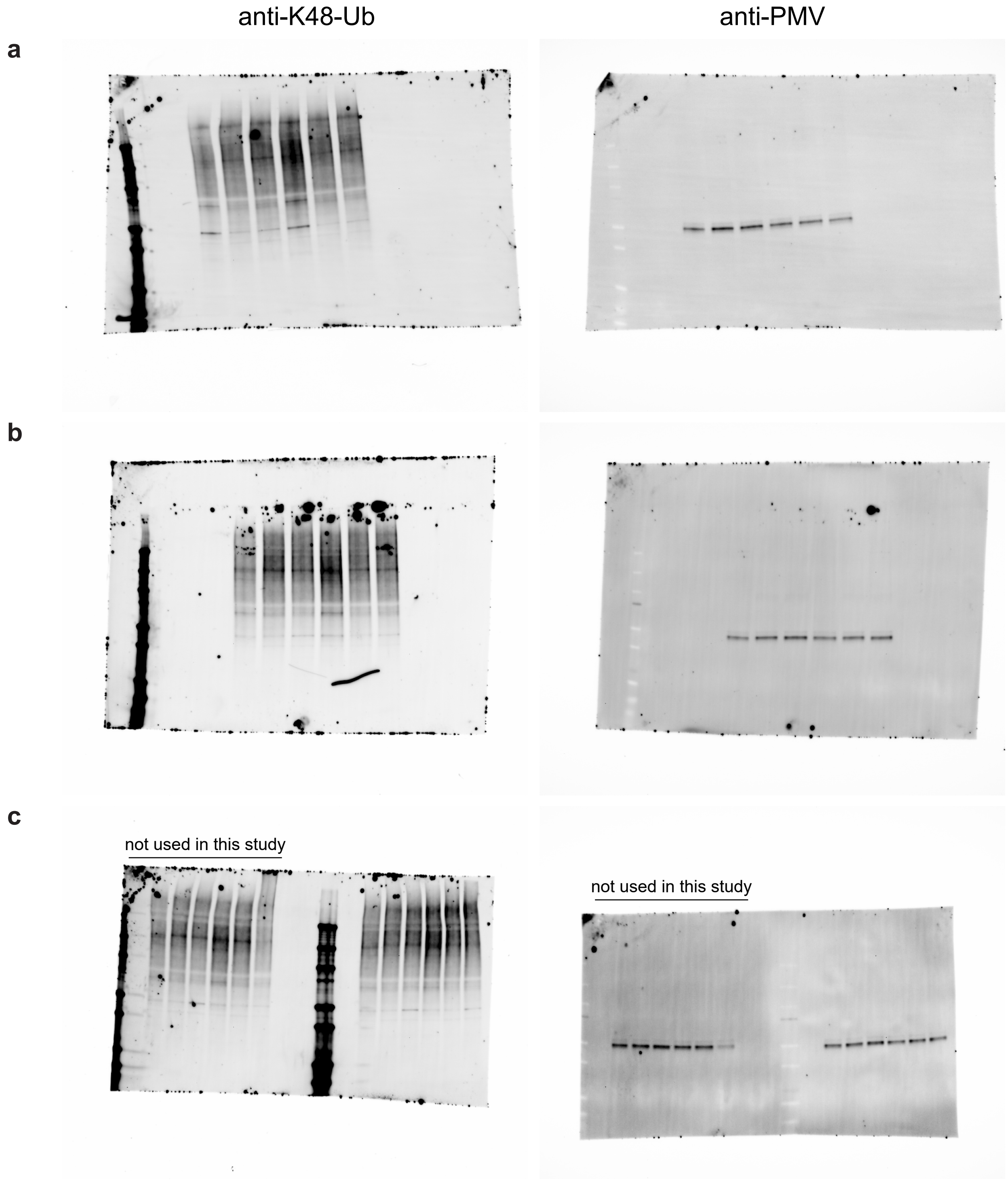


**Figure 3- source data 1. Uncropped western blots used for ImageJ quantification.** Dd2 trophozoites were treated with 10 μM CQ, 25 nM DHA, 10 μM CQ + 25 nM DHA for 6 h. Parasite lysates were subjected to western blot and probed with anti-K48-linked ubiquitin (K48-Ub) antibodies or anti-plasmepsin V (PMV) antibodies. Blots from three independent replicates are shown.

**Figure 4- figure supplement 1**

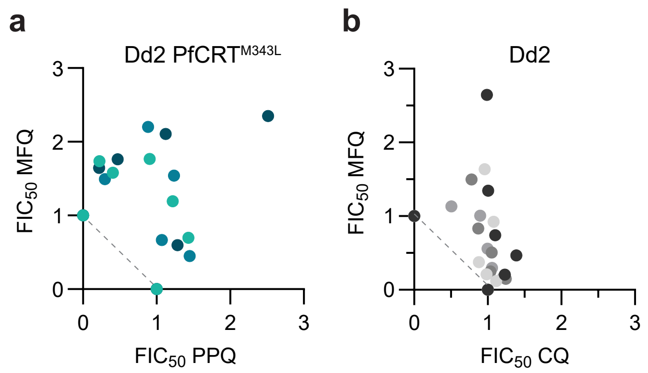


**Figure 4- figure supplement 1. Additional MFQ isobolograms.** Trophozoite stage isobolograms were performed on (**a**) Dd2 PfCRT$^{M343L}$ and (**b**) Dd2 parasites to determine drug-drug interactions of using the following fixed ratios: 1:0, 4:1, 2:1, 1:1, 1:2, 1:4, 0:1. Shown are fractional IC50 values from 3 independent replicates.

**Figure 4- figure supplement 2**

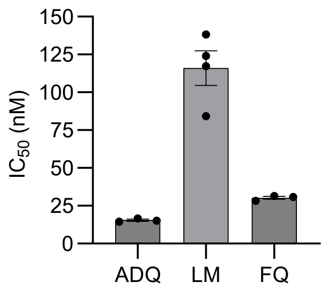


**Figure 4- figure supplement 2. ADQ, LM, and FQ IC50 values.** Shown are mean Dd2 trophozoite stage IC50 values ± SEM obtained as part of independent isobologram replicates (see Figure 4d, e, and g).

### Figure 3 and 4- table supplement 1

| Parasite | Dd2 | | | | Dd2 | | | | Dd2 PfCRT$^{Dd2}$ | | | Dd2 PfCRT$^{3D7}$ | | | Dd2 | | | | |
|---|---|---|---|---|---|---|---|---|---|---|---|---|---|---|---|---|---|---|---|
| Treatment Time | 5.5 h | | | | 1 h | | | | 5.5 h | | | 5.5 h | | | 5.5 h | | | | |
| Treatment | – | 10 µM CQ | 150 nM PPQ | 150 nM MFQ | – | 10 µM CQ | 10 µM PPQ | 10 µM MFQ | – | 150 nM CQ | 10 µM CQ | – | 150 nM CQ | 10 µM CQ | – | 10 µM CQ | 150 nM FQ | 150 nM ADQ | 500 nM LM |
| Mean intensity | 13.44 | 1.31 | 6.63 | 13.42 | 11.41 | 4.66 | 6.32 | 9.10 | 8.37 | 5.48 | 1.76 | 9.27 | 6.09 | 4.01 | 10.90 | 0.71 | 3.51 | 4.88 | 10.28 |

**Figure 3 and 4- table supplement 1. Mean parasite intensity for Ac-H-FluNox experiments.** Parasite line, treatment duration, and antimalarial treatment, and mean Ac-H-FluNox are indicated.
